# Enhancer regulation for induced *WNT3A* expression during neuronal regeneration

**DOI:** 10.1101/861153

**Authors:** Chu-Yuan Chang, Jui-Hung Hung, Ching-Chih Wu, Min-Zong Liang, Pei-Yuan Huang, Joye Li, Hong-I Chen, Shaw-Fang Yet, Ka Shing Fung, Cheng-Fu Kao, Linyi Chen

## Abstract

The treatment of traumatic brain injury (TBI) is limited by a lack of knowledge about the mechanisms underlying neuronal regeneration. WNT family members have been implicated in neurogenesis and aberrant WNT signaling has been associated with neurodegenerative diseases. The current study compared the expression of WNT genes during regeneration of injured cortical neurons. Recombinant WNT3A showed positive effect in promoting neuronal regeneration via *in vitro* and *in vivo* TBI models. Intranasal administration of WNT3A protein to TBI mice increased NeuN^+^ cells compared to control mice as well as retained motor function based on behavior analysis. Since TBI is known to reprogram the epigenome, chromatin immunoprecipitation-sequencing of histone H3K27ac and H3K4me3 was performed to address the transcriptional regulation of *WNT3A* during neuronal regeneration. We predicted, characterized and proposed that a histone H3K4me1-marked enhancer may undergo topological transformation to regulate the *WNT3A* gene expression.

## 1. Introduction

Traumatic brain injury (TBI) is a major cause of morbidity and lifelong disability, making it a critical healthcare concern worldwide. Globally, an estimated 69 million individuals per year suffer from TBI, and according to the Centers for Disease Control and Prevention, it accounts for over 30% of injury-related death in the United States (Roozenbeek *et al*. 2013; Fehily & Fitzgerald 2016; Dewan *et al*. 2018). There is currently no effective treatments can stimulate regeneration, partly due to the heterogeneity of pathophysiology, severity and outcomes, so much of the damage/dysfunction in TBI patients becomes permanent (McGuire *et al*. 2018). Among over 200 registered ongoing or completed for TBI, administration of chemical compounds (e.g., sertraline, amantadine, minocycline), transplantation of autologous bone marrow-derived cells, and other interventions may partially rehabilitate impaired consciousness and/or improve neurocognitive dysfunction. Nevertheless, a large number of these trials have been terminated (*ClinicalTrials.gov*), and the rising medical burden of TBI highlights the pressing need to develop better treatments for brain injury.

Poor regeneration of the injured central nervous system (CNS) is thought to largely result from inhibitory gliosis and an aging-related decline of intrinsic regenerative capacity (Yiu & He 2006; Giger *et al*. 2010; Shen *et al*. 2009). Inhibiting the non-permissive environment alone has not been a successful strategy to promote axonal regeneration in injured brains (Fawcett & Asher 1999; Lee *et al*. 2009; Sun & He 2010). We reason that modulation of a subset of regeneration-associated genes (RAGs) may be needed to stimulate intrinsic neuronal regeneration programs. In the case of dorsal root ganglions, when injured, remodeling of the epigenome leads to differential expression of RAGs, such as *Gfpt1* (glutamine fructose-6-phosphate transaminase 1) and *Fxyd5* (FXYD domain containing ion transport regulator 5) (Chandran *et al*. 2016). However, the RAGs that function in brain neurons appear to be distinct from those in the peripheral nervous system or spinal cord (Bareyre *et al*. 2011; Bomze *et al*. 2001; Seijffers *et al*. 2007; Yang & Yang 2012). This tissue specificity may arise from the regulation of RAGs through distal enhancers, which are functional regulatory DNA elements that drive cell-type- and tissue-specific patterns of gene expression (Calo & Wysocka 2013; Li *et al*. 2016). The abundance of specific chromatin signatures, such as enriched histone H3 lysine 27 acetylation (H3K27ac) or a high ratio of histone H3 lysine 4 mono-methylation to tri-methylation (H3K4me1:H3K4me3), suggests that enhancers far outnumber protein-coding genes in the human genome, reflecting their critical role in transcriptional control (Lam *et al*. 2014; Kim *et al*. 2015). Given this important role in transcriptional activation, identifying enhancers that regulate RAGs may be a useful first step in developing strategies to activate regeneration programs. Databases and computational methods for the identification of enhancers have been built based on analyses of sequences, chromatin states or combinations of features (Bu *et al*. 2017; He *et al*. 2017; Singh *et al*. 2018). However, accurate prediction of distal enhancers with high resolution remains challenging because of genetic variation across different cell types, tissues and species (Lim *et al*. 2018).

WNT signaling is an evolutionarily conserved pathway that regulates axis specification of the neural plate, neural tube morphogenesis, dendrite and axon development, and synaptic plasticity (Mulligan & Cheyette 2012; Munji *et al*. 2011; Salinas & Zou 2008). Aberrant WNT signaling has previously been implicated in Alzheimer’s disease and Parkinson’s disease (Bayod *et al*. 2015; Marzo *et al*. 2016; Zhang *et al*. 2016). Studies also reported differential expression of WNT proteins being implicated in the responses to spinal cord injury yet little is known regarding the precise regulation of *WNT* genes (Fernandez-Martos *et al*. 2011; Gonzalez-Fernandez *et al*. 2014; Lambert *et al*. 2016; Strand *et al*. 2016). In this study, we determined the expression of *WNT* genes during regeneration of injured cortical neurons. *In vitro* and *in vivo* studies demonstrated the regenerative potential of WNT3A upon traumatic brain injury. Mechanistically, we identified, characterized and proposed an enhancer regulation on regeneration-induced expression of *WNT3A* gene.

## 2. Materials and methods

### 2.1 Animal experiments and ethics approval

All experiments were executed in accordance with the guidelines of the Laboratory Animal Center of National Tsing Hua University (NTHU), and protocols were approved by the NTHU Institutional Animal Care and Use Committee. The study was not pre-registered. No randomization and blinding were performed in this study. No statistical method was used to determine sample size. Sample size was arbitrarily set within 10.

### 2.2 Reagents

Powder of Minimum Essential Medium (MEM) and Dulbecco’s Modified Eagle Medium (DMEM), fetal bovine serum (FBS), horse serum (HS), penicillin-streptomycin (PS), B-27™ supplement, L-glutamine (L-Gln), Antibiotic-Antimycotic (AA), TRIzol reagent, Lipofectamine 2000, Alexa Fluor 488 (Cat#A21206; RRID: AB_2534069), 555 (Cat#A21422; RRID: 141822) IgG secondary antibodies and 4’, 6’-Diamidino-2-phenylindole (DAPI; Cat#D1306; RRID: AB_2629482) were purchased from Invitrogen (Carlsbad, CA). Neurobasal^®^ medium, Eagle’s basal medium (BME), and N-2 Supplement were purchased from Gibco (Grand Island, NY). Poly-L-lysine (PLL), glutamate, Cytosine-β-D-arabinofuranoside (AraC), bovine serum albumin (BSA), sucrose, formaldehyde solution, chloroform, phenol solution, and IWR-1 were from Sigma-Aldrich (Saint Louis, MO). WNT3A recombinant protein (Cat#315-20) was purchased from PeproTech (Rocky Hill, NJ). WNT8A (Cat#8419-WN-010/CF) and WNT9B (Cat#3669-WN-0.25/CF) were from R&D Systems (Minneapolis, MN). RNase A and *Power* SYBR Green Master Mix were purchased from Thermo Fisher Scientific (Waltham, MA). T7EI (Cat#M0302) was purchased from New England BioLabs (Ipswich, MA). Anti-Tau (Cat#sc-5587; RRID: AB_661639) and anti-MAP2 (Cat#sc-56561; RRID: AB_2138164) antibodies were purchased from Santa Cruz Biotechnology (Dallas, TX). Anti-TUJ1 antibodies (Cat#802001/RRID: AB_2564645 and Cat#801202/RRID: AB_10063408) were purchased from BioLegend (San Diego, CA). Anti-H3K4me3 antibody (Cat#39159; RRID: AB_2615077) was purchased from Active Motif (Carlsbad, CA). Anti-H3K27ac (Cat#ab4729; RRID: AB_2118291) and anti-H3K4me1 (Cat#ab8895; RRID: AB_306847) antibodies were from Abcam (Cambridge, UK). Anti-GFAP (Cat#GTX10877), anti-NeuN (Cat#GTX30773; RRID: AB_1949456) antibodies and rabbit IgG (Cat#GTX26702; RRID: AB_378876) were from GeneTex (Irvine, CA) or Novus (Cat#NB300-141; RRID: AB_10001722; Centennial, CO) for cryosection staining.

### 2.3 *In vitro* culture of primary neurons, injury/regeneration assay, neurite tracking, immunofluorescence staining and quantification of neurite re-growth

Sprague-Dawley (SD) rats were purchased from BioLASCO Taiwan Co., Ltd. Pregnant female rats (P12-14; body weight: 300-350g) were anaesthetized with 15 mg Zoletil^®^ 50 (Virbac, Carros, France)/rat via intraperitoneal injection. Buprenorphrine (0.02 mg/kg body-weight/rat) was intraperitoneally injected to mitigate the pain. Abdominal incision was performed to expose the E18 embryos. Following embryos take-away, female rats were scarified via cutting through the diagram. Primary cortical neurons were dissociated from dissected cortices of rat embryos (E18) as previously described (Chen *et al*. 2015; Shih *et al*. 2013). Cells were seeded on PLL-coated plates. On day *in vitro* (DIV) 0, primary neurons were cultured in MEM supplemented with 5% FBS, 5% HS, and 0.5 mg/ml PS under 5% CO_2_ condition. Culture medium was changed to Neurobasal^®^ medium containing 25 μM glutamate, 2% B-27™ supplement, 0.5 mM L-Gln, and 50 units/ml AA on DIV1. AraC (10 μM) was added to neurons on DIV2 to inhibit proliferation of glial cells. On DIV3, medium was changed to Neurobasal^®^ medium containing 2% B-27™ supplement, 0.5 mM L-Gln and 50 units/ml AA. After DIV3, half the medium was exchanged with fresh Neurobasal/Glutamine medium every 3 days.

For the injury assay, primary neurons were seeded on PLL-coated wells. On DIV8, neurons were injured by scraping with a p20 tip to generate two injured gaps in each well (Fig. 1a). For samples with WNT recombinant protein treatments, cortical neurons were pre-treated with WNT3A (50 ng/ml), WNT8A (100 ng/ml), WNT9B (100 ng/ml), or phosphate buffered saline (PBS) control 1 h prior to injury. For treatment with WNT inhibitor IWR-1, cortical neurons were treated with IWR-1 (10 µM) or DMSO control (1%) immediately after injury. Neurite re-growth was monitored from iDIV8 to iDIV11 in the same field-of-view by using the “Mark and Find” positioning function of AxioVision software (Zeiss). Live images were captured on a Carl Zeiss Observer Z1 microscope. The average width of injured gaps was calculated from measurements made at twelve positions in each gap. The remaining gap distance was measured at the indicated time points. The percentage of gap closure was calculated as 100% × (1 - remaining gap distance/original gap distance). Up to six injured gaps were quantified per condition in at least three independent experiments.

**Figure 1.**
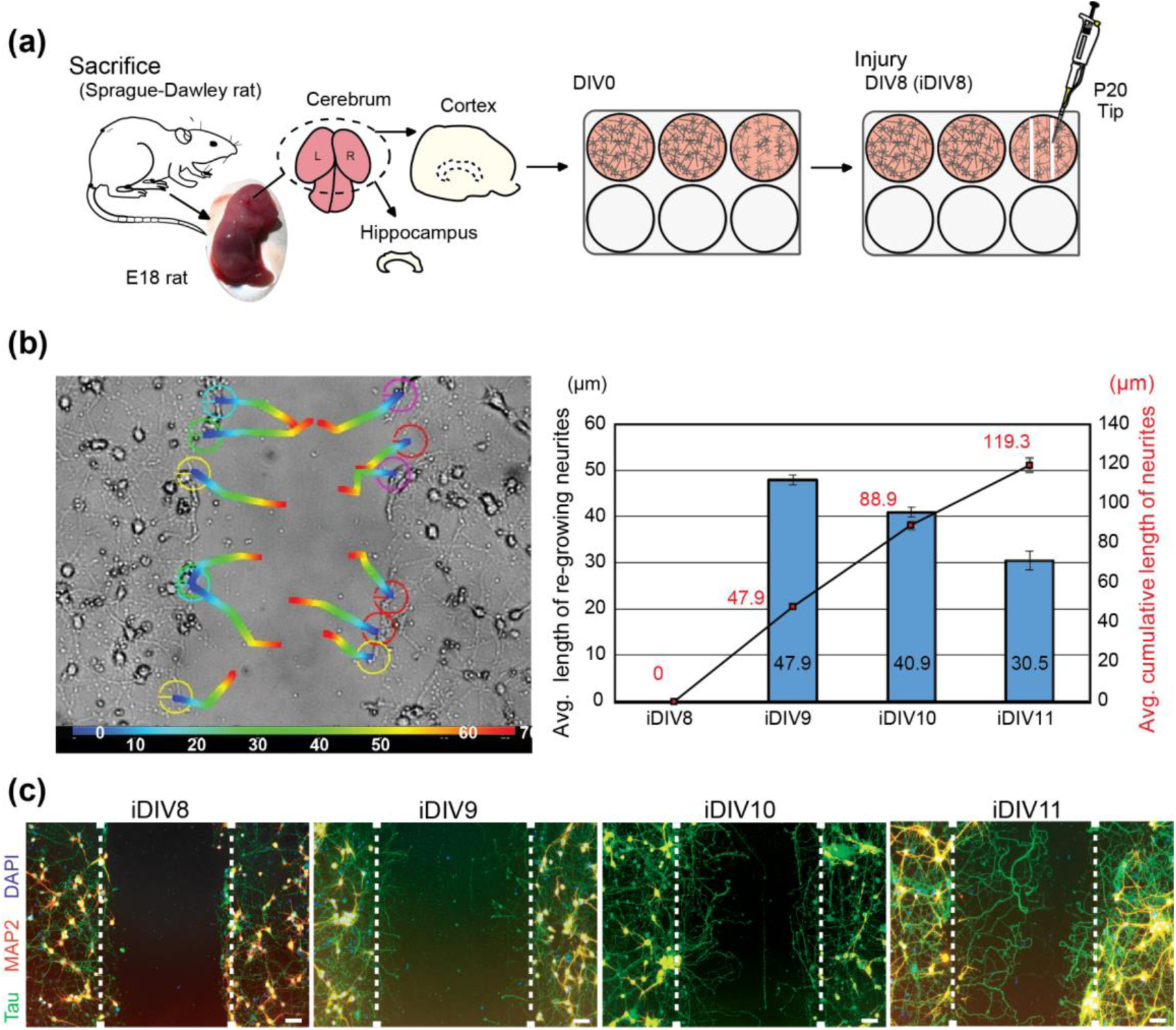
Neurite re-growth of primary cortical neurons in an *in vitro* TBI model. (a) Schematic flow chart of experimental design. Neurite re-growth was quantified to represent neuronal regeneration. (b) Trajectory of single neurite re-growth was tracked from 0 h (blue) to 72 h (red) after injury using tracking function of AxioVision software (upper). Quantitative results of average length of re-growing neurite (blue bars) and average cumulative length (line graph) are shown from iDIV8 to iDIV11 (bottom). (n = 3, track 12 neurites/experiment.) (c) Cortical neurons were fixed on iDIV8 to iDIV11 and were subjected to immunofluorescence staining with anti-Tau (green, axonal marker), anti-MAP2 (red, dendritic marker) antibodies and DAPI (blue, nuclei). Merged images are shown. White dashed lines indicate the borders of the injured gap. Scale bar, 50 µm.

To track re-growth of individual neurons, the tracking function of AxioVision software (Zeiss) was used on live images from the injury assay. From iDIV8 to iDIV11, the re-growth of 12 single neurites per field was tracked in three injured gaps in three independent experiments. The tips of regenerating neurites were located at 24-h intervals after injury, and the length of extension was measured for each day. The average length of neurite re-growth per day and the average cumulative re-growth (0-72h) were calculated.

To better visualize neurites, neurons were fixed, blocked and incubated with specific primary antibodies. Anti-Tau, anti-MAP2, and anti-TUJ1 antibodies were used at the dilution of 1:200, 1:300, and 1:500 respectively. Fluorescence images were taken using a Carl Zeiss Observer Z1 microscope.

### 2.4 ChIP-seq, RNA-seq, and superarray qPCR

For Chromatin immunoprecipitation-sequencing (ChIP-seq), ChIP was performed as previously described (Dahl & Collas 2007). In brief, an appropriate number of control and DIV8-injured cortical neurons were fixed on DIV9 and DIV10 with 1% (v/v) formaldehyde; 1.25 M glycine was added to quench the cross-linking reaction. Lysates were collected and sonicated to produce DNA fragments of 200-500 bp length. Sheared chromatin was pulled down by H3K4me3 or H3K27ac antibodies. DNA was purified with a MinElute^®^ PCR Purification Kit (Qiagen). DNA libraries were prepared by using the TruSeq^TM^ ChIP Library Prep Kit (Illumina) and were sequenced on an Illumina HiSeq 2500 System as 2 × 100-bp paired ends, following the manufacturer’s instructions; all sequencing was performed at the Next Generation Sequencing (NGS) High Throughput Genomics Core (Academia Sinica). FASTQ files were generated by the BclToFastq Pipeline Software (Illumina) followed by adaptor trimming by using PEAT (Li *et al*. 2015) and alignment to the rat genome (Rnor 6.0/rn6) using Bowtie v1.1.1 (Langmead *et al*. 2009) with the following parameters (-n 1 --best -l 100 -strata -a -M 1). SAM-to-BAM file format conversion was performed using SAMtools standard software for further data processing (Li *et al*. 2009). To display sequencing coverage of the genome, BAM files were converted to BigWig files using Bedtools2 v2.19.1 (Quinlan & Hall 2010), and then BigWig files were uploaded onto JBrowse Genome Browser (Skinner *et al*. 2009). Peak calling was performed by MACS v1.4.2 (https://github.com/taoliu/MACS/) to identify histone ChIP-seq enrichment over input with the following parameters (--nomodel –shiftsize 73 –pvalue=0.01) (Zhang *et al*. 2008).

For RNA-sequencing (RNA-seq), total RNAs were isolated from cortical neurons by TRIzol reagent. Ribosomal RNA-depleted RNA-seq libraries were prepared using a Ribo-Zero+TruSeq Stranded mRNA Library Prep Kit (Illumina) and processed for 2 × 150-bp paired-end sequencing on an Illumina HiSeq 2500 System at the NGS High Throughput Genomics Core (Academia Sinica). FASTQ files were generated by the BclToFastq Pipeline Software (Illumina) and adaptor trimming was accomplished with PEAT (Li *et al*. 2015). The RNA-seq reads were mapped to rn6 using STAR v2.4.1 with the following arguments (--outSAMstrandField intronMotif --chimSegmentMin 15 -- chimJunctionOverhangMin 15) (Dobin *et al*. 2013). Expression levels of RNA transcripts were analyzed by using HTseq v0.6.1 with the following parameters (htseq-count -s no -m intersection-nonempty -a 0 -t gene) (Anders *et al*. 2015). The TMM (trimmed mean of M-value) normalization (Robinson *et al*. 2010) across samples was then applied to each gene, and the values were used to calculate differential gene expression.

For superarray analysis, total RNA was extracted from cortical neurons. Isolated RNA was processed and used in a WNT Signaling Pathway-focused PCR Array (SuperArray Bioscience Corporation). Differential gene expression levels were calculated.

### 2.5 Data analysis

#### Aggregation plot

To visualize the overall enrichment and signal distribution of H3K4me3 across gene promoters in each sample, we aggregated normalized ChIP-seq signals across a ± 4 kb region flanking the transcription start sites (TSSs) of 88 WNT-related genes. Normalized ChIP-seq signals were calculated as reads per million (RPM).

#### ChromHMM for enhancer prediction

To predict enhancers, ChromHMM was used for chromatin-state discovery on the basis of chromatin modification patterns by model training and analysis (Ernst & Kellis 2012). ChIP-seq datasets utilized for learning chromatin-state models were collected from our H3K4me3, H3K27ac, KLF4 and KLF7 ChIP-seq profile of control and injured cortical neurons (on DIV9 and DIV10), and Gene Expression Omnibus (GEO) accession numbers: GSE41217 (H3K9me3 modification in hippocampal neurons), GSE22878 (RNA polymerase II occupancy in cortical neurons), GSE64703 (Sox10 occupancy in spinal cord), GSE63103 (H3K27ac modification in Schwann cells), and GSE64971 (H3K27ac modification in injured sciatic nerves). Processed BED files containing genomic information were used as the input for chromatin-state annotation. Genome-wide chromatin was segmented into 10 states. To identify potential enhancers, modeled emission parameter and positional enrichment profiles were referenced. High enrichment of H3K27ac and relatively low H3K4me3 and RNA Polymerase II at distal regions to the TSSs indicated chromatin regions that were annotated as State2, and these regions were selected as putative enhancers. Positional information of the segments in BED files were uploaded onto JBrowse Genome Browser for visualization.

To predict enhancers of *WNT* genes, putative enhancers within the region ranging from -1 Mb to +100 kb to the TSS of *WNT3A* gene (chr10: 45,514,600-46,690,800) were separated into 10 domains, e1-e10. To analyze overall enrichment and signal distribution of H3K27ac across each enhancer in each sample, we combined RPM-normalized ChIP-seq density across a ±2 kb region flanking the central base pair of enhancers within e1-e10 individual enhancer domains (shown in Fig. 5e).

#### Long-range chromatin interaction prediction

To predict possible interaction between putative enhancers, publicly available Hi-C datasets of human hippocampal tissues (sample GSM2322543 from series GSE87112) (Schmitt *et al*. 2016) was analyzed in the 3D-genome Interaction Viewer and database (3DIV) (Yang *et al*. 2018). The “Interaction visualization” function was used with query gene names of *WNT3A* and *MPRIP*, the gene located within the e5 enhancer region, to visualize chromatin interactions around *WNT3A* gene and predicted enhancer e5-e7 region, respectively.

#### ENCODE datasets processing

ChIP-seq datasets were collected from the ENCODE Project Consortium (Consortium 2012) and visualized on UCSC Genome Browser with the alignment to mouse genome assembly (GRCm38/mm10). H3K27ac and H3K4me1 ChIP-seq data of mouse forebrain tissue were extracted from ENCSR428OEK (H3K27ac; E16.5), ENCSR094TTT (H3K27ac; P0), ENCSR141ZQF (H3K4me1; E16.5), ENCSR465PLB (H3K4me1; P0). CCCTC-binding factor (CTCF) ChIP-seq data of mouse forebrain tissue was extracted from ENCSR677HXC (CTCF; P0). Multiz alignments and conservation track are shown. Homologous loci of putative enhancers e1-e10 between rat and mouse genome were aligned and indicated in the shadowed regions in the track view (shown in Fig. 6c).

### 2.6 ChIP-qPCR assays and eRNA quantification

For histone ChIPs, DIV8-injured or uninjured control cells were cross-linked on DIV9 and DIV10 with 0.8% formaldehyde. Chromatin was sonicated to generate 200-500 bp fragments for subsequent immunoprecipitation of histone-DNA complexes using anti-H3K4me3, anti-H3K4me1, or anti-rabbit IgG antibody-conjugated protein A. Final DNA purification was carried out by RNase A and Proteinase K treatments, followed by phenol/chloroform DNA extraction. The immunoprecipitated DNA and input control were analyzed by qPCR using *Power* SYBR Green Master Mix and an ABI StepOnePlus Real-Time PCR System. Specific primers were designed to amplify *WNT3A*, *WNT9B* and *WNT10A* promoter regions and putative enhancers e5, e7 and e10.

For qPCR quantification, total RNA or ChIP DNA were prepared from primary cortical neurons at indicated time points and analyzed by qPCR using *Power* SYBR Green Master Mix and an ABI StepOnePlus Real-Time PCR System. For gene expression analysis, total RNA from neurons was isolated by TRIzol reagent following the manufacturer’s instructions and processed with a High-Capacity cDNA Reverse Transcription Kit (Cat#4368814; Applied Biosystems) before qPCR. Relative gene expression was calculated by the ddCt method, normalized to *GAPDH*. For the ChIP assay, immunoprecipitated DNA and input control were prepared as described above. The enrichment of histone modification at promoters or enhancers was determined as percentage of input (% input) = 100 × 2^{Ct(ChIP) – [Ct(input) - log2(input dilution factor)]}. Input dilution factor = 1/(fraction of the input chromatin saved).

For eRNA quantification, total RNA was isolated for reverse transcription polymerase chain reaction (RT-PCR) assays. eRNAs of interest were amplified by specific primers and analyzed by DNA electrophoresis. Band intensities were quantified by GelPro31 software and normalized to *GAPDH*.

### 2.7 Organotypic brain slice culture

Brains isolated from E18 rats were washed with HBSS medium and embedded in 2.5% low-melting agarose gel. The brain was then soaked in the artificial cerebrospinal fluid (ACSF: 125 mM NaCl, 5 mM KCl, 1.25 mM NaH_2_PO_4_, 1 mM MgSO_4_, 2 mM CaCl_2_, 25 mM NaHCO_3_ and 20 mM glucose; pH 7.4) pre-pumped with 95% O_2_/5% CO_2_ and sectioned by a Leica microtome VT100. Coronal tissue sections with a thickness of 350 µm were collected and injured with a scalpel as indicated. Sliced tissues were cultured on PLL-coated inserts in 66% BME medium supplemented with 25% HBSS, 5% FBS, 1% N-2 Supplement, 1% P/S, and 0.66% (wt/vol) D-(+)-glucose. For WNT3A treatment, WNT3A recombinant protein (50 ng/ml) was added daily, along with fresh culture medium. AraC (5 μM) was administered from DIV0 to DIV2 to limit the proliferation of glial cells.

For immunofluorescence staining of brain slices, injured brain slices were fixed with 4% PFA for 2 h at room temperature on iDIV4. The insert membrane with the brain slice attached was trimmed and tissues were subjected to immunostaining of anti-TUJ1 and anti-GFAP antibodies at dilutions of 1:750 and 1:500 respectively overnight at 4°C and1:1000 secondary antibodies overnight at 4°C. DAPI was used to stain the nuclei at room temperature for 2 h. The brain slices were mounted and images were taken with a Carl Zeiss LSM780 confocal microscope. For quantification of regenerating injured brain slice, the area that contained of regenerating neurites and the length of injured tissue border were measured by Zeiss ZEN 2.3 lite Imaging Software. The length of regenerating neurites was calculated as (area/length).

### 2.8 Controlled cortical injury (CCI) model

Eight-to-ten-week old C57BL/6J male mice were purchased from National Laboratory Animal Center, Taiwan, and subjected to TBI by using the pneumatic CCI equipment (Electric Cortical Contusion Impactor, Custom Design & Fabrication, Inc., the United States). Inhalation anesthesia was performed with 3-5% isoflurane and maintained anesthetic of the mice with 1.5-3% isoflurane. Buprenorphine (0.2 mg/100 g body-weight/mouse) was intraperitoneally injected to mitigate the pain. Upon anesthesia, the mouse was fastened on the stereotaxic frame. Craniectomy was then performed to remove the skull in a shape of 3-mm-diameter circle over left frontoparietal cortex (−0.5 mm anteroposterior and +0.5 mm mediolateral to bregma). After exposing the dura matter, the brain was impacted by pneumatically operated CCI device at the velocity of 3 m/s, reaching 1 mm in depth, affecting a 2-fmm-diameter circular area, and with 500-ms dwell time to generate mild TBI (Yu *et al*. 2009). After CCI-induced brain injury and suturing, PBS or 50 ng WNT3A was intranasally delivered to mice daily for consecutively four days. On 4 dpi, the TBI mice were anesthetized and perfused with saline and 4% paraformaldehyde. The brain tissues were then isolated, immersed in 10%, 15%, 20% sucrose followed by embedded in Shandon Cryomatrix embedding resin (Cat#6769006; Thermo Scientific) and sliced into 10-μm sections by cryostat microtome (Leica CM3050 S). The cryosections were incubated in antigen retrieval solution at 70°C for 20 min and then incubated in 1% BSA for 2 h. After BSA blocking, the cryosections were subjected to immunofluorescence staining with anti-TUJ1 and anti-GFAP antibodies, followed by incubating with fluoresce-conjugated secondary antibodies. DAPI is used to stain the nuclei at room temperature for 10 min. Finally, the cryosections were mounted by using 90% glycerol. Fluorescence images of brain cryosections were imaged by Zeiss Confocal LSM800 and analyzed by Zeiss ZEN 2.3 lite Imaging Software. The intensity of GFAP around the impact site (indicated as “proximal”), at the ipsilateral hemisphere distal to the impact site (indicated as “distal”) and at the contralateral uninjured hemisphere (indicated as “uninjured”), was quantified separately by calculating the average GFAP intensity of 4 defined squared region of interest (ROI) in each brain region. Relative intensity of GFAP was defined as the average intensity of proximal regions divided by the average intensity of distal or uninjured regions (Fig. 4b, middle panel). The number of NeuN^+^ cells was defined as the average NeuN^+^ cells counted in the 4 squared ROI at the proximal regions around the impact site. The area of each ROI is 0.05 mm^2^. The numbers of animal used are specified in the Results section.

### 2.9 Behavior assay for functional recovery

Asymmetry use of forelimb was measured to represent functional recovery of damaged brain region. To this end, the cylinder test was conducted on 1 day before injury (-1 dpi, day post injury), and 1-3 dpi, during 7-9 pm to quantify the asymmetry use of forelimbs. The timeline of the experimental design is depicted below.

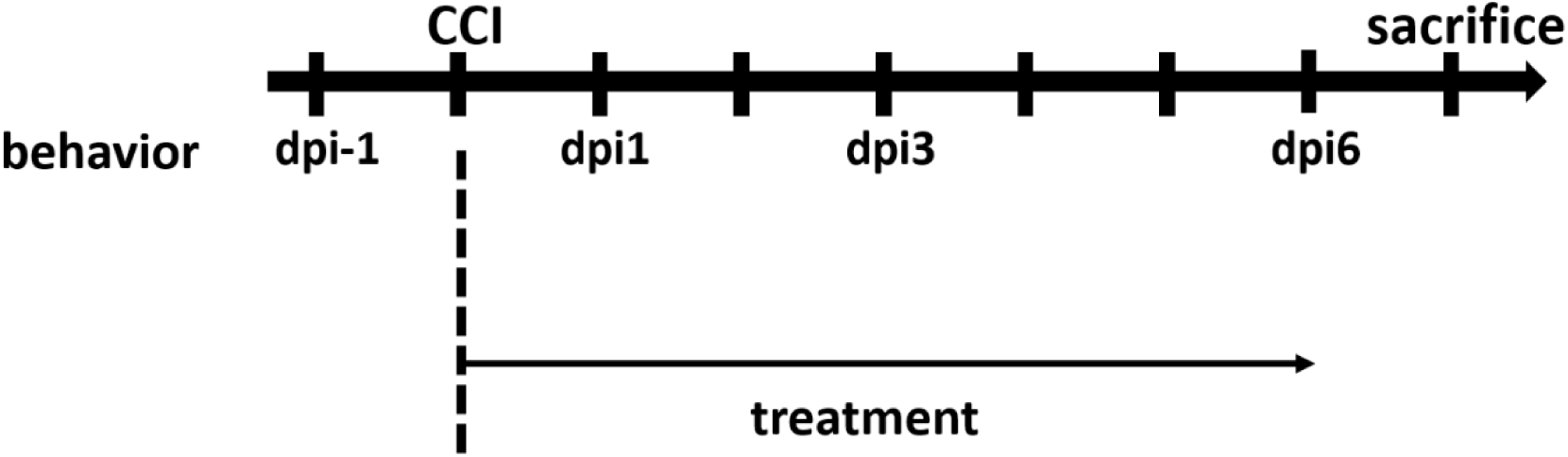

The protocol of cylinder test is modified according to Hua, Y *et al*.’s (Hua *et al*. 2002). Briefly, C57BL/6 mice were put in a transparent cylinder of plexiglass and the behavior was recorded. The number of times that the forelimbs contact on the cylinder wall was calculated in 5 minutes. Valid forelimb-wall contacts were considered for further data quantification if the forelimbs were raised over the shoulder of the mice. Data of cylinder test are quantified by calculating the percentage of the unimpaired left forelimb use (ipsilateral limb to the CCI site) which was calculated by following parameter: (1) the frequency of the mice using its left forelimb first (U, unimpaired) (2) the frequency of the mice using its right forelimb first (I, impaired) (3) the frequency of the mice using both forelimbs (B, both). The percentage of the unimpaired limb use was calculated as [U / (U+I+B)], indicated as “% left forelimb use”. To minimize bias, data collected from behavior experiments of sham group animals demonstrating slight brain injury or CCI animals failed to be injured at expected brain regions were excluded, according to immunostaining of the cryosections. Among available behavior recordings, only data where forelimb contact equals to or more than three times in the given duration were included.

### 2.10 Statistical analysis

All results are expressed as mean ± S.E.M. or box-and-whisker plot from at least three independent experiments. Statistical significance (*) of parametric data is defined as *P* ≤ 0.05 in a paired Student’s *t*-test, unless otherwise noted. For behavior tests, statistical significance (*) is defined as *P* ≤ 0.05 in non-parametric Mann-Whitney U-test. Microsoft^®^ Excel^®^ 2016 MSO (16.0.4849.1000) 64 bits was used for statistical analysis.

## 3. Results

### 3.1 Identification of WNT genes as RAGs for axonal regeneration of cortical neurons

To evaluate neuronal regeneration, rat cortical neurons were cultured *in vitro*. On DIV8, primary cortical neurons were injured by scraping lines through the culture with a p20 tip (Fig. 1a). It was estimated that 10% neurons were injured via this approach. An anti-mitotic reagent, AraC, was added to the culture medium to inhibit proliferation of glial cells (Brusehafer *et al*. 2014; Liu *et al*. 2013). Regenerating neurites were tracked for 72 h after injury. Regeneration of injured cortical neurons was most prominent within 48 h (iDIV10) after the injury (Fig. 1b). Immunofluorescence staining using anti-microtubule-associated protein 2 (MAP2, marker for dendrites) and anti-Tau (axonal marker) antibodies demonstrated that the re-growing neurites were mostly Tau^+^ axons (Fig. 1c).

To identify candidate RAGs during regeneration of injured cortical neurons, our previous RNA-seq data found that approximately 43% of total up-regulated 28,055 genes are between 1 to 2 fold during regeneration [12,170 genes (43.4%) for iDIV9/DIV9; 12,303 genes (43.9%) for iDIV10/DIV10]. Importantly, among these up-regulated genes, there are enriched protein-coding genes that are directly or indirectly WNT signaling-related. For example, leucine-rich repeat containing G protein-coupled receptor 5 (*LGR5*) was 1.5-to-1.6-fold up-regulated on iDIV9 and iDIV10 compared to DIV9 and DIV10 while lymphoid enhancer-binding factor 1 (*LEF1*) was 1.2-to-1.3-fold up-regulated. Along this line, RNA-seq data revealed clusters of *WNT* genes being up-regulated in iDIV9/DIV9 (15 out of 19 *WNT*s) and iDIV10/DIV10 (14 out of 19 *WNT*s) cortical neurons (Fig. 2a-b). There were 12 out of 19 *WNT* genes were up-regulated in 1-2 fold in response to injury on both iDIV9 and iDIV10. A pathway-focused superarray analysis of 88 *WNTs* and WNT target genes showed that 48 genes (54.5%) were up-regulated. Among them, *WNT3*, *WNT3A, WNT4, WNT5A, WNT5B, WNT6, WNT7A, WNT8A, WNT8B, WNT9A, WNT9B* and *WNT10B* expression were up-regulated (fold change > 1.5) during regeneration (Fig. 2c). Increased expression of *WNT3A, WNT8A, WNT9B* on iDIV10 compared to DIV10 was validated by qPCR, with the expression of *WNT3A* and *WNT9B* showing significant increase (Fig. 2d). Although the fold increase of *WNT* genes is not high, local concentration of secreted WNTs in the brain tissue under tight control of blood-brain-barrier (BBB) may be sufficient to trigger cellular response.

**Figure 2.**
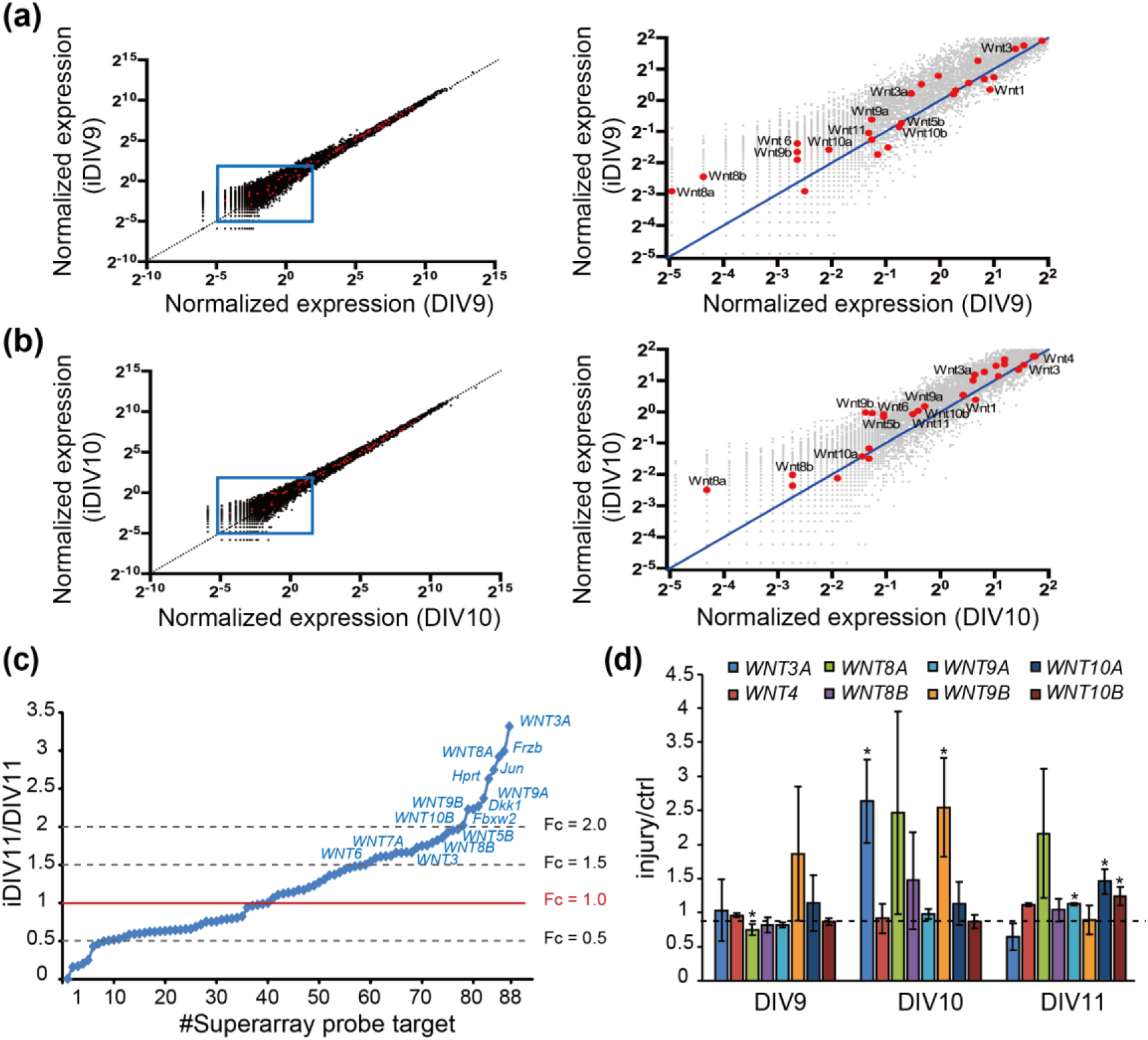
Identification of regeneration-associated genes via RNA-seq and superarray analysis. (a, b) Scatter plots showing normalized gene expression levels on DIV9 versus iDIV9 and iDIV10 versus DIV10 respectively, as determined from RNA-seq. Data points representing TMM-normalized expression of *WNT* genes are marked as red dots. The blue lines indicate equal expression between uninjured control and injured samples. Right panels are the close-up view of the boxed regions on the left panels. (c) Differential expression profile of WNT-related genes by superarray assay is shown. Genes with an expression fold-change above 1.5 in response to injury are denoted. (d) Analysis of differential expression of *WNT*s by qPCR during neuronal regeneration. Data are presented as mean ± SEM from at least three independent experiments (*WNT3A*, *8A*: n = 4; *WNT9B*, *10A*: n = 5). \**P* ≤ 0.05 (paired Student’s *t*-test).

### 3.2 WNT3A recombinant protein promotes neuronal regeneration and functional recovery

To address the effect of WNT proteins on the regeneration of injured cortical neurons directly, WNT3A, WNT8A, WNT9B, and WNT10A recombinant proteins were added. The percentage of relative regeneration was determined as average distance of neurite re-growth compared to the initial width of the injury gap. As shown in Fig. 3a-b, the regeneration of injured cortical neurons was improved by 20-25% in the presence of WNT3A compared to without WNT3A by 72 h (iDIV11). The addition of WNT8A, WNT9B, and WNT10A also enhanced regeneration of injured cortical neurons, though not to the same extent as WNT3A did (Fig. 3a).

**Figure 3.**
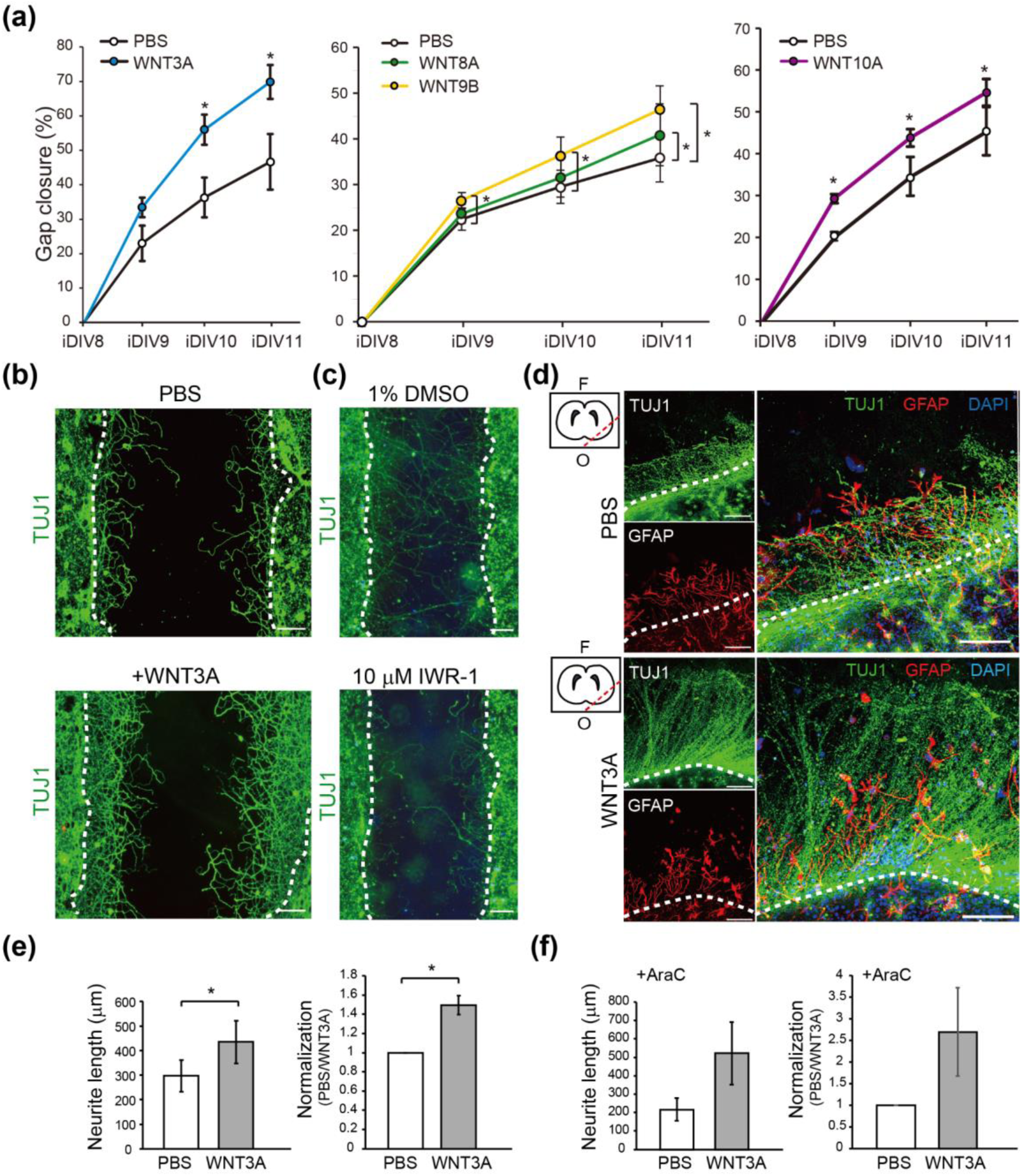
Recombinant WNT3A promotes regeneration of injured cortical neurons and brain tissues. (a) Cortical neurons were pre-treated with PBS, WNT3A, WNT8A, WNT9B or WNT10A recombinant proteins prior to injury on DIV8. The percentages of gap closure were calculated from iDIV8 to iDIV11. Data are presented as mean ± SEM (n = 4 for WNT3A and WNT9B experiments; n = 5 for WNT10A experiment). \**P* ≤ 0.05 (paired Student’s *t*-test). (b) Cortical neurons with or without WNT3A (50 ng/ml) pre-treatment were fixed and subjected to immunofluorescence staining with anti-TUJ1 antibody (neurites) on iDIV11. (c) Cortical neurons were treated with 1% DMSO or IWR-1 (10 µM) prior to injury on DIV8. Neurons were subjected to immunofluorescence staining with anti-TUJ1 antibody and the representative images are shown. Dashed lines in (b) and (c) indicate the borders of the injured gap. Scale bar, 100 µm. (d) The effect of WNT3A on neuronal regeneration was assessed using organotypic brain slice culture. Brain slices were injured at the olfactory tubercle on DIV0, as indicated by the red dashed line in the upper left atlas map, and cultured with or without WNT3A (50 ng/ml) in AraC-containing medium. Brain tissues were fixed on iDIV4 and subjected to immunofluorescence staining with anti-TUJ1 (green) and anti-GFAP (cyan, glia) antibodies, and DAPI (blue). The border of injured sites is depicted by white dashed lines, and remaining brain tissue is underneath the lines in each panel. F: frontal lobe; O: occipital lobe. Scale bar, 100 µm. (e, f) The length of regenerating neurites from the brain slices treated with (e) PBS control or WNT3A and (f) AraC+PBS control or AraC+WNT3A were calculated. Data on the right panels in (e) and (f) is normalized to the PBS control, indicated as fold change. Data are presented as mean ± SEM (n = 4). \**P* ≤ 0.05 (paired Student’s *t*-test).

**Figure 4.**
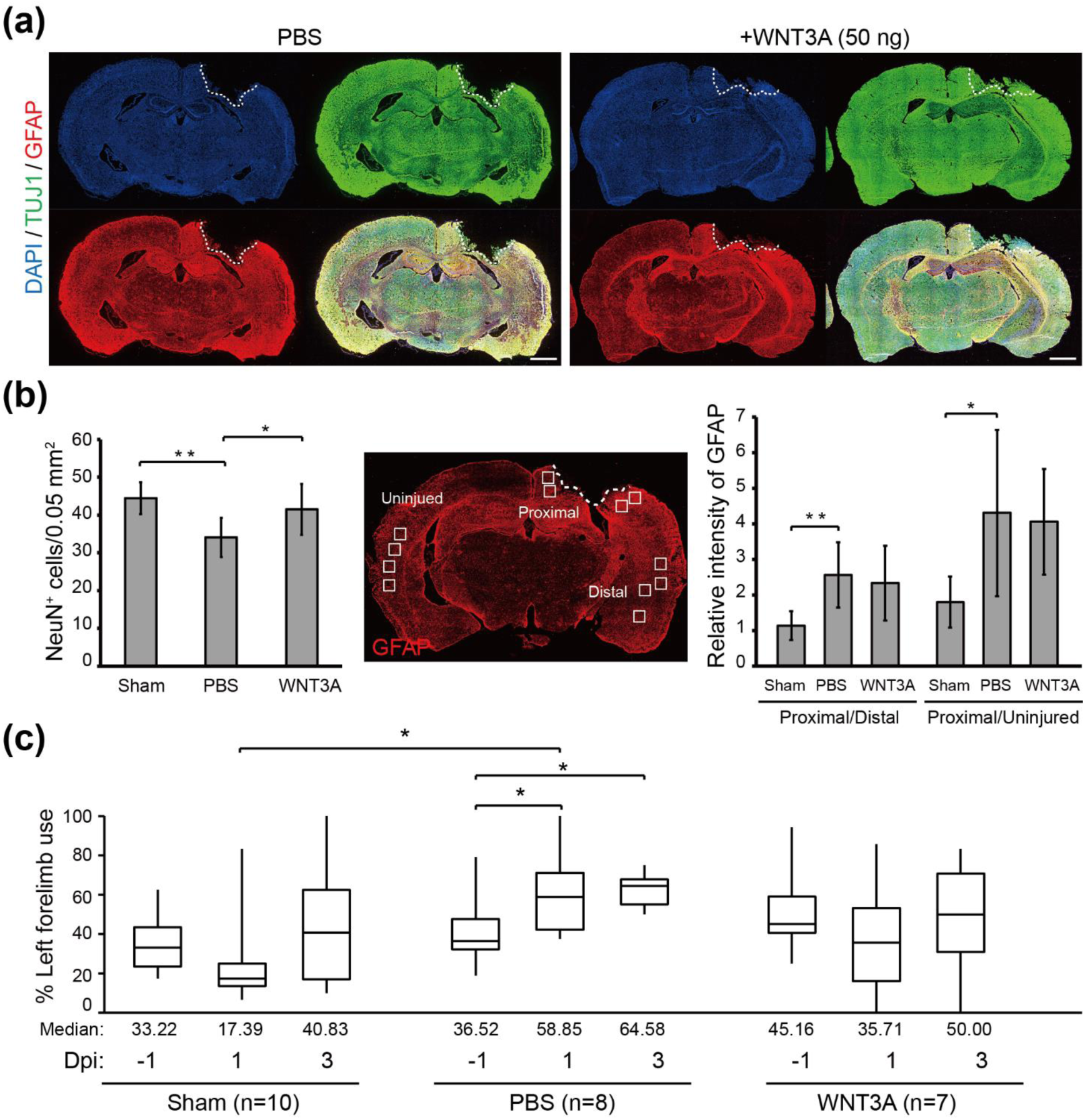
Recombinant WNT3A promotes regeneration of injured brain tissue and preserves motor function of CCI mice. (a) The effect of WNT3A on neuronal regeneration was assessed using *in vivo* CCI mouse model. Cortical impact on motor and somatosensory cortex was generated and the mice were given PBS or 50 ng WNT3A intranasally during 0-3 dpi. Injured brain tissues were fixed, cryosectioned on 4 dpi and subjected to immunofluorescence staining with anti-TUJ1 (green), anti-GFAP (red) antibodies and DAPI (blue). White dashed lines depict the border of CCI sites. Scale bar, 1 mm. (b) The number of NeuN^+^ cells (left) and the relative intensity of GFAP (right) of CCI the cryosections from sham group (n = 7) and PBS-treated (n = 8) or WNT3A-treated (n = 6) CCI animals were quantified as described in the Materials and Methods. The middle panel of GFAP-immunostained cryosection defines 0.05 mm^2^-ROIs within white boxes. Dashed lines in indicated the borders of the impact site. Data are presented as mean ± SEM. \**P* ≤ 0.05, \*\**P* ≤ 0.01 (paired Student’s *t*-test). (c) The box plots of mice forelimb placing capacity in the left (ipsilateral unimpaired) forelimb before and after CCI for sham control (n = 10), PBS control (n = 8), and WNT3A treatment (n = 7) (-1 to 3 dpi). The boundaries of the boxes closest and farthest to the zero point indicate the 25^th^ and the 75^th^ percentiles, respectively. The black bars within the boxes mark the median and the values are presented below the plot. The limit of whiskers above and below the boxes mark the maximal and minimal values respectively. \**P* ≤ 0.05 (Mann-Whitney U-test).

**Figure 5.**
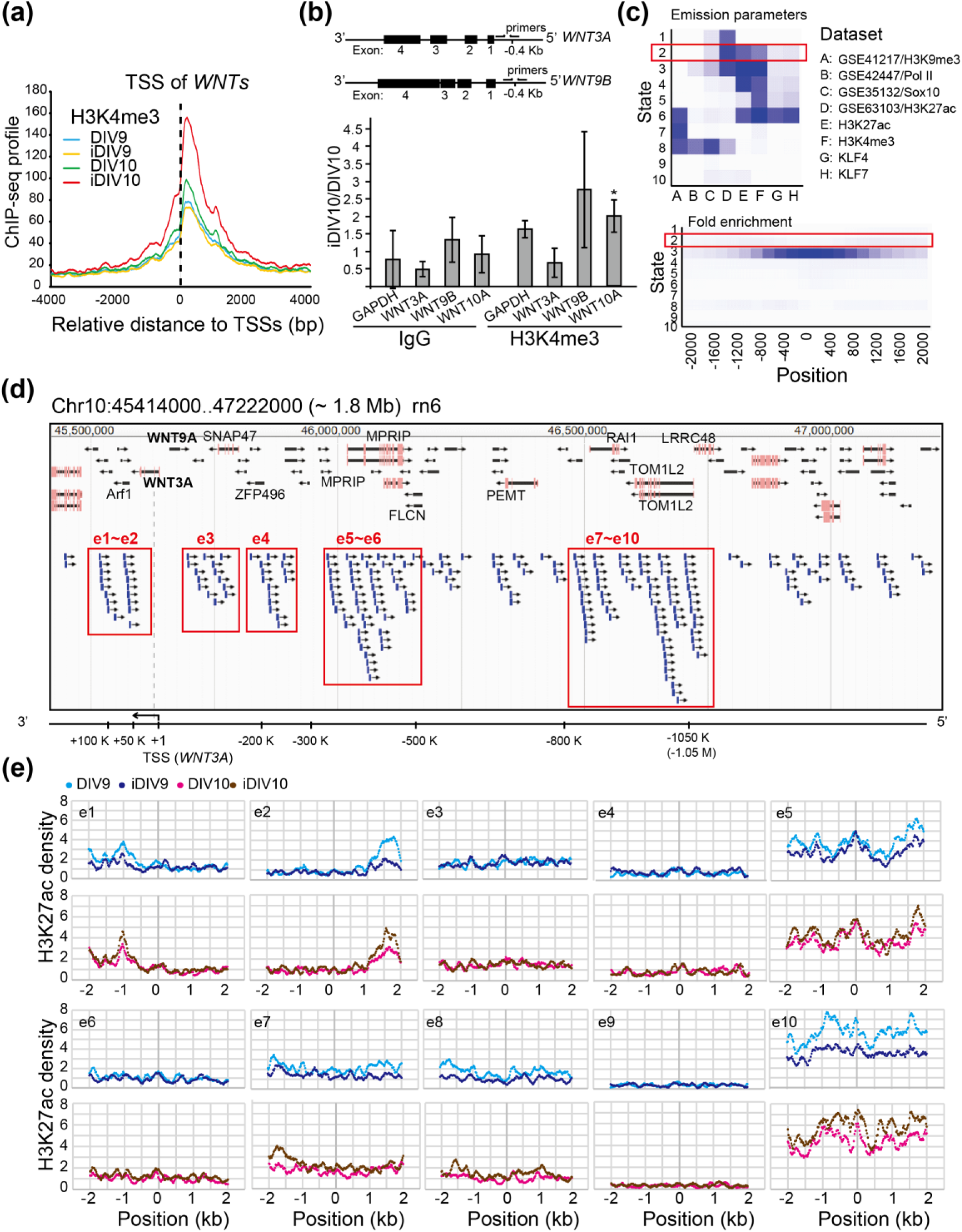
Prediction of putative enhancers of *WNT3A* gene during neuronal regeneration. (a) Aggregation of normalized H3K4me3 signal density profiles of 88 WNT-related genes across the ± 4 kb promoter regions. H3K4me3 signals across control and injured samples are indicated by colored lines. (b) Upper: Schematic diagrams of gene tracks for rat *WNT3A* and *WNT9B*. Targeted loci for specific primers in the promoters used for ChIP assays are shown. Bottom: The fold-change of H3K4me3 at the promoters of *WNT3A*, *WNT9B*, or *WNT10A* in cortical neurons comparing DIV10 and iDIV10 were analyzed by ChIP-qPCR. The *GAPDH* promoter was used as a negative control. Data were normalized to IgG and then to DIV10 controls, indicated as fold change. Data are presented as mean ± SEM from three independent experiments. \**P* ≤ 0.05 (paired Student’s *t*-test). (c) Functional enrichment of chromatin states in rat genome performed by ChromHMM. Upper: Heatmap of the model parameter with chromatin states numbered in the emission order. The columns refer to relative enrichment for the indicated annotation in corresponding chromatin states. ChIP-seq data from four GEO datasets as well as data from this study were used to train the model. Bottom: Heatmap of the positional enrichment of annotated chromatin state. The genomic feature of the State2 elements is indicated in red boxes. (d) Snapshot of JBrowser Genome Browser demonstrating the region across 1.8 Mb flanking the *WNT3A* TSS of the rat genome (RCSC 6.0/rn6). Ten predicted genomic regions (e1-e10, indicated in boxed area) may be enhancers for the *WNT3A* gene based on the enrichment of clustered State2 elements assigned by ChromHMM. A diagram of *WNT3A* gene region is shown at the bottom. (e) Aggregation of normalized H3K27ac ChIP-seq profiles across the ± 2 kb central base pair of within each putative enhancers region (e2-e10) for control (DIV9, DIV10) and injured (iDIV9, iDIV10) samples.

**Figure 6.**
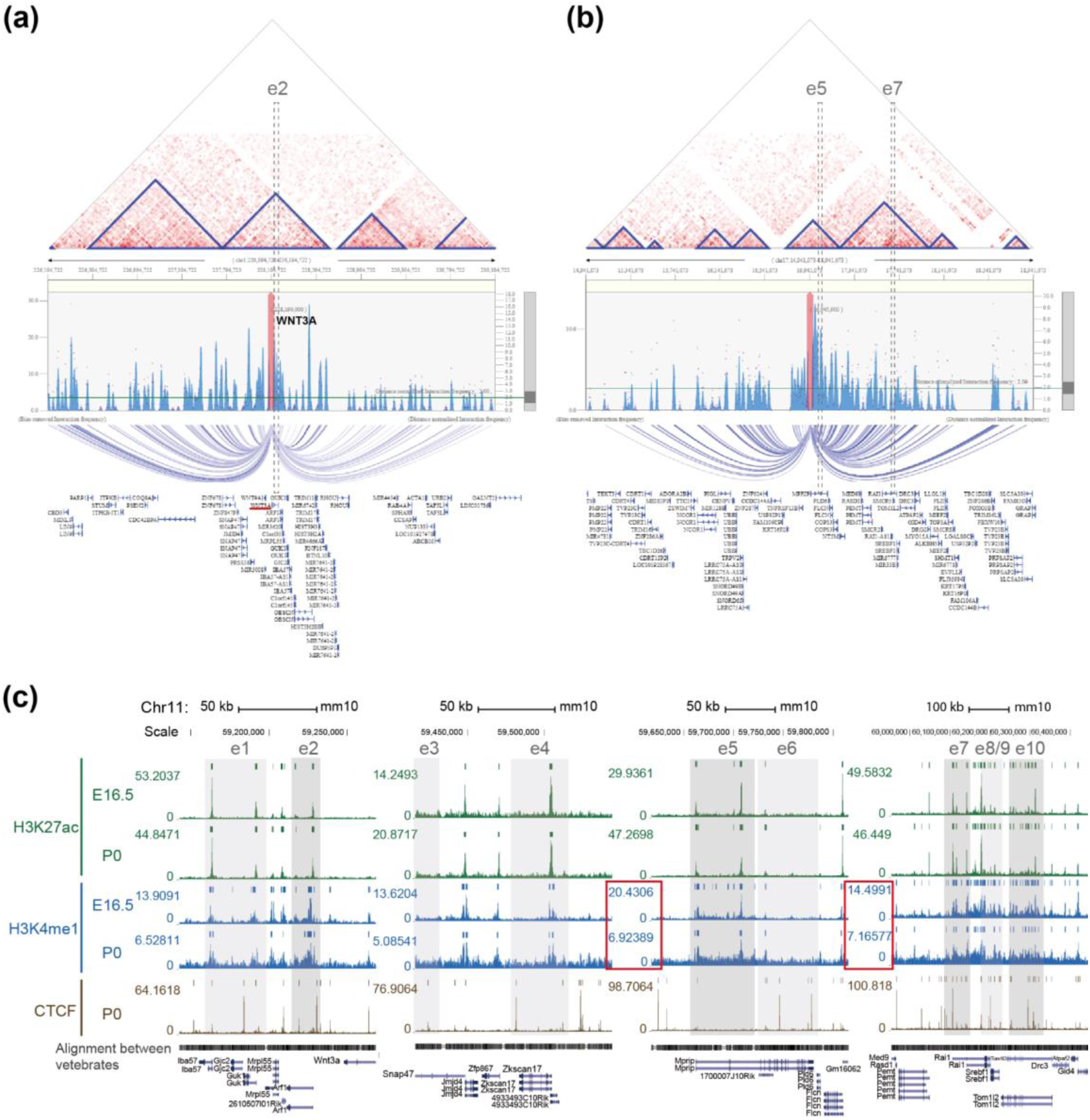
Chromatin interactions and H3K4me1modifications at the e5 and e7 enhancers. (a, b) Visualization of identified chromatin interactions around (a) e2/*WNT3A* and (b) e5/e7 regions. Blue triangles in the interaction heatmaps indicate TADs that form 3D structural compartment of the genome. Interaction frequency graphs, consisting of the blue bars and magenta dots, represents bias-removed and distance-normalized interaction frequencies respectively. Arc-diagrams anchored by indicated loci (*WNT3A* and *MPRIP* genes, indicated by magenta bars in the frequency graphs,) are defined by indicated threshold (distance-normalized interaction frequencies ≥ 2) and shown. RefSeq annotations are denoted at the bottom of each panels. (c) Homologous loci of predicted enhancer domains aligned with the mouse genome assembly were enriched with enhancer marks (H3K4me1 and H3K27ac). CTCF binding profile is also shown. ChIP-seq data of the indicated samples were collected from the ENCODE database and visualized in UCSC Genome Browser. Colored bars above peaks in each track indicate confidence of enrichment for the corresponding genomic feature. Putative enhancers are shown in gray shadowed areas. e2, e5, e7, e10 are highlighted by darker gray shadows. H3K4me1 signals at e5-e6 and e7-e10 are indicated by the red boxed region. Genomic sequence alignment among vertebrates and gene tracks are depicted below ChIP-seq profiles.

Thus, the WNT3A-dependent regeneration was further examined by treating injured cortical neurons with IWR-1, a WNT3A inhibitor, and neuronal regeneration was inhibited (Fig. 3c). To assess the effect of WNT3A in a multicellular three-dimensional (3D) culture, organotypic brain slice culture was used. Brain slices were injured in the olfactory tubercle, followed by WNT3A or PBS treatment. In order to reduce injury-induced glial cell proliferation, which has been shown to hinder neurite re-growth, effect of combining WNT3A+AraC or PBS+AraC was also compared. As shown in Fig. 3e-f, injured brain slice treated with WNT3A significantly increased the length of regenerating neurites compared to PBS control. To show the *in vivo* effect of WNT3A administration on neuronal regeneration, we adapted one of the TBI models, controlled cortical impact (CCI) injury mice model. CCI model mimics post-TBI neuro-pathophysiology, resembling cognitive or behavioral deficits seen in human TBI patients (Yu *et al*. 2009; Xiong *et al*. 2013; O’Connor *et al*. 2011). C57BL/6J mice were injured via CCI at 3 m/s in speed and 1 mm in depth to generate mild TBI, followed by daily intranasal administration of 50 ng WNT3A recombinant protein for consecutive four days. As demonstrated in Fig. 4a, administration of WNT3A robustly enhanced regeneration of brain tissue on 4 dpi, characterized by better healing of the injured area compared to vehicle treatment. Within the regenerated tissue, there were increased NeuN^+^ (neuronal marker) cells for WNT3A-treated mice compared to PBS-treated control group (Fig. 4b, left panel). Upon injury, increased glial fibrillary acidic protein (GFAP) reflects injury-induced proliferation of glial cells. GFAP^+^ signal was compared between proximal regions to the injury site, ipsilateral distal regions to the injury site, and contralateral uninjured regions. The relative intensity of GFAP at the proximal region over distal region (proximal/distal) was higher in PBS-treated group compared to that of sham-treated group, whereas there was no significant difference between sham-treated group and WNT3A-treated group (Fig. 4b, middle and right panels). There was also no difference between PBS-treated and WNT3A-treated groups. This result suggests that WNT3A administration does not reduce injury-induced proliferation of glial cells. Functional recovery of damaged motor and somatosensory brain regions was assessed by cylinder test conducted on 1 day before injury (-1 dpi), 1, and 3 dpi to quantify the asymmetry use of forelimb. If left brain was injured, motor function of the right forelimb would be impaired. A preferred usage of left forelimb (asymmetry use) would be observed. As revealed in Fig. 4c, sham-treated mice exhibited 10% variation of asymmetry score from 1-3 dpi compared to that of -1 dpi. PBS administration increased 20-25% asymmetry score from 1-3 dpi compared to that of -1 dpi, reflecting obvious brain injury. WNT3A administration reduced asymmetry score 10% on 1 dpi and increased 5% on 3 dpi. These results suggest that WNT3A may protect neurons or promote neuronal regeneration during the first three days after TBI.

### 3.3 Putative enhancer region for WNT3A expression

To investigate the mechanism by which injury-induced *WNT*s expression is regulated, we had performed ChIP-seq analysis of histone H3K4me3 and H3K27ac modifications. The normalized H3K4me3 ChIP-seq profiles of WNT-related genes were aggregated across ± 4 kb promoter regions with TSSs as the anchors (Fig. 5a). The aggregated ChIP-seq signals aligned to the proximal promoters of WNT-related genes were increased up to 1.6-fold on iDIV10 compared to DIV10. ChIP-qPCR analyses for H3K4me3 at promoters of *WNT*s were then performed. As shown in Fig. 5b, active H3K4me3 mark at the gene promoter of *WNT10A* was increased, whereas those for *WNT3A* and *WNT9B* were not. If the transcriptional regulation of *WNT3A* and *WNT9B* was not through promoter, it might be regulated by a distal enhancer.

To test this suggestion, a number of computational tools have been employed to predict candidate enhancers during regeneration of injured cortical neurons. For example, ChromHMM, an approach for chromatin-state discovery and characterization (Ernst & Kellis 2012), identified candidate enhancer regions based on extracting general chromatin features of enhancers. We classified the rat genome into 10 states, according to histone codes and the occupancy of transcription factors (Fig. 5c). To do so, we utilized our reference datasets of H3K4me3, H3K27ac, Krüppel-like factor 4 (KLF4) and Krüppel-like factor 7 (KLF7) ChIP-seq profiles, as well as published ChIP-seq datasets of H3K9me3, RNA polymerase II (RNAPII), Sox10, and H3K27ac from the GEO (GSE41217, GSE22878, GSE64703, GSE63103, and GSE64971) of multiple cell types in the nervous system for training chromatin-state models. Sites with highly enriched H3K27ac marks (active enhancers) and no occupancy of RNAPII (Fig. 5c, upper panel) that are distal to the TSS (Fig. 5c, bottom panel) in genomic regions, annotated as State2, are likely to represent enhancer regions (Fig. 5c). We then defined ten genomic regions within a 1.8-Mb interval flanking the *WNT3A* TSS in the rat genome (RCSC 6.0/rn6). These regions, e1 to e10, represented putative enhancer domains that may regulate the expression of *WNT3A* genes. Each predicted domain contains a high density of State2 genomic elements assigned by ChromHMM (Fig. 5d). Notably, this 1.8-Mb region also includes the *WNT9A* gene, and it is possible that *WNT3A* and *WNT9A* share common enhancers for coordinated transcriptional control. To screen the enhancer regions that are active during neuronal regeneration, H3K27ac ChIP-seq signals were evaluated. As shown in Fig. 5e, H3K27ac signals mapped to each putative enhancer domain were aggregated. The ChIP-seq signal density for H3K27ac was elevated at the e2, e5, e7, e10 enhancer regions in iDIV10 samples compared to DIV10. The H3K27ac modification in e1 and e8 were slightly increased, whereas no obvious change could be observed in e3, e4, e6, and e9 regions. In contrast, H3K27ac modification in e1-e10 was generally decreased or remained unchanged throughout the enhancer regions on iDIV9 compared to DIV9. Thus, the candidate enhancer for induced *WNT3A* expression may lie within e2, 5, 7 and 10.

While the predicted enhancers for the *WNT3A* gene reside within the region from +100 kb to -1 Mb to the TSS (e1-e10), it is reasonable to speculate that the regulation of *WNT3A* expression is governed by spatial chromatin interaction between functional genomic elements, for instance, promoter-enhancer interaction. To investigate possible interactions between *WNT3A* promoter and the putative enhancers, significant interactions around *WNT3A* gene promoter were identified by the 3D Interaction Viewer and database (3DIV) online tool (https://www.kobic.kr/3div/). 3DIV collected publicly available Hi-C data in human cell/tissue types and provides normalized chromatin frequencies as well as browsing visualization tool for scientists to further interpret genome-wide chromatin interactions (Yang *et al*. 2018). As shown in Fig. 6a, several topologically associating domains (TADs) were identified within an interaction range of 2 Mb around *WNT3A* in the genome of human hippocampal tissue (Schmitt *et al*. 2016). The genomic region homologous to the e2 enhancer in the rat genome resides within one of the TADs that comprises *WNT3A* gene and presents relatively high interaction frequency to *WNT3A*. Moreover, according to the GeneHancer database, an integrated human enhancer database that infers enhancer-gene associations (Fishilevich *et al*. 2017), also suggests a high likelihood score of this region as potential elite promoter/enhancer for *WNT3A* gene (GeneHancer score: 2.5; Gene Association Score: 10.6; Total score: 25.53). The relative position of e2 to *WNT3A* is similar between human and rat, with the TSS distance of +79.5 kb and +57.8 kb to the e2 region, respectively. Through mapping the homologous loci of rat e5-e7 region to the human genome, a serendipitous finding revealed an evolutionary shift of e5-e7 region to a different chromosome in human (chromosome 17), different from e2 region and *WNT3A* gene (chromosome 1). Nonetheless, the homologous e5 and e7 regions reside within two separated TADs and the data suggests an interaction between e5 and e7 (Fig. 6b). This finding uncovers a possibility of long-range inter-chromosomal regulation of *WNT3A* gene in human.

To determine whether these putative enhancer regions contain enhancer signature, we compared several enhancer-specific histone marks. While there is currently no publicly accessible ChIP-seq database for rat brain, we compared ChIP-seq datasets for H3K27ac and H3K4me1 enhancer marks in mouse forebrain that are available in the ENCODE database (https://www.encodeproject.org/) (Rosenbloom *et al*. 2013). Homologous enhancer regions between rat and mouse were identified, and the annotations of putative enhancer domains in the rat genome were converted to the mouse genome (Casper *et al*. 2018; Kent *et al*. 2002). ENCODE annotation data for the homologous regions is shown in Fig. 6c. Overlapping enhancer marks were found within homologous enhancer domains, with a clear decrease of the H3K4me1 mark in these regions from E16.5 to postnatal day 0 (P0), according to ChIP-seq. For example, normalized signal of H3K4me1 mark in the e5-e6 regions reduced around 3 folds (from 20.4 to 6.9) during development while around 2 folds (from 7.1 to 14.5) decrease in the e7 regions (Fig. 6c, indicated in the red boxes); however, the level of H3K27ac mark remained relatively constant over the same time period. This differential decrease of histone modification suggests that the H3K4me1 may prime and activate enhancers during development and thus could be a cue for RAG regulation during neuronal regeneration. The existence of CTCF sites surrounding e1-e2, e4, e5-e6, and e7-e10 regions probably reflects a functional role of the CTCF architectural protein in bridging enhancer-promoter interacting topology (Ong & Corces 2014; Ren *et al*. 2017) (Fig. 6c). To examine the role of the H3K4me1 modification at putative enhancers of *WNT3A*, ChIP assays of H3K4me1 modification at the e5, e7, e10 were performed before and after injury. As shown in Fig. 7a-c, the H3K4me1 modification at e5 sub-regions, e5-1, e5-2, and e5-3, significantly increased on iDIV9 and iDIV10 compared to uninjured controls. Similarly, the H3K4me1 mark at e7 sub-region, e7-3, significantly increased during regeneration (Fig. 7a, d-e). H3K4me1 modification at e10 was not detectable via qPCR. Notably, given that recent studies have linked the expression of non-coding enhancer RNAs (eRNAs) to functional enhancers (Ding *et al*. 2018; Zhu *et al*. 2013; Meng & Bartholomew 2018), the expression of cognate eRNA derived from the e5 and e7 were also examined. As shown in Fig. 7f-g, the relative expression of e5-3, e7-4, e7-5 eRNA significantly increased on iDIV9. Expression of *MPRIP* and *RAI1* genes, located within the e5 and e7 regions respectively, did not change during regeneration (data not shown). These results suggest that the H3K4me1-modulated e5/e7 region may be an active enhancer for *WNT3A* during regeneration.

**Figure 7.**
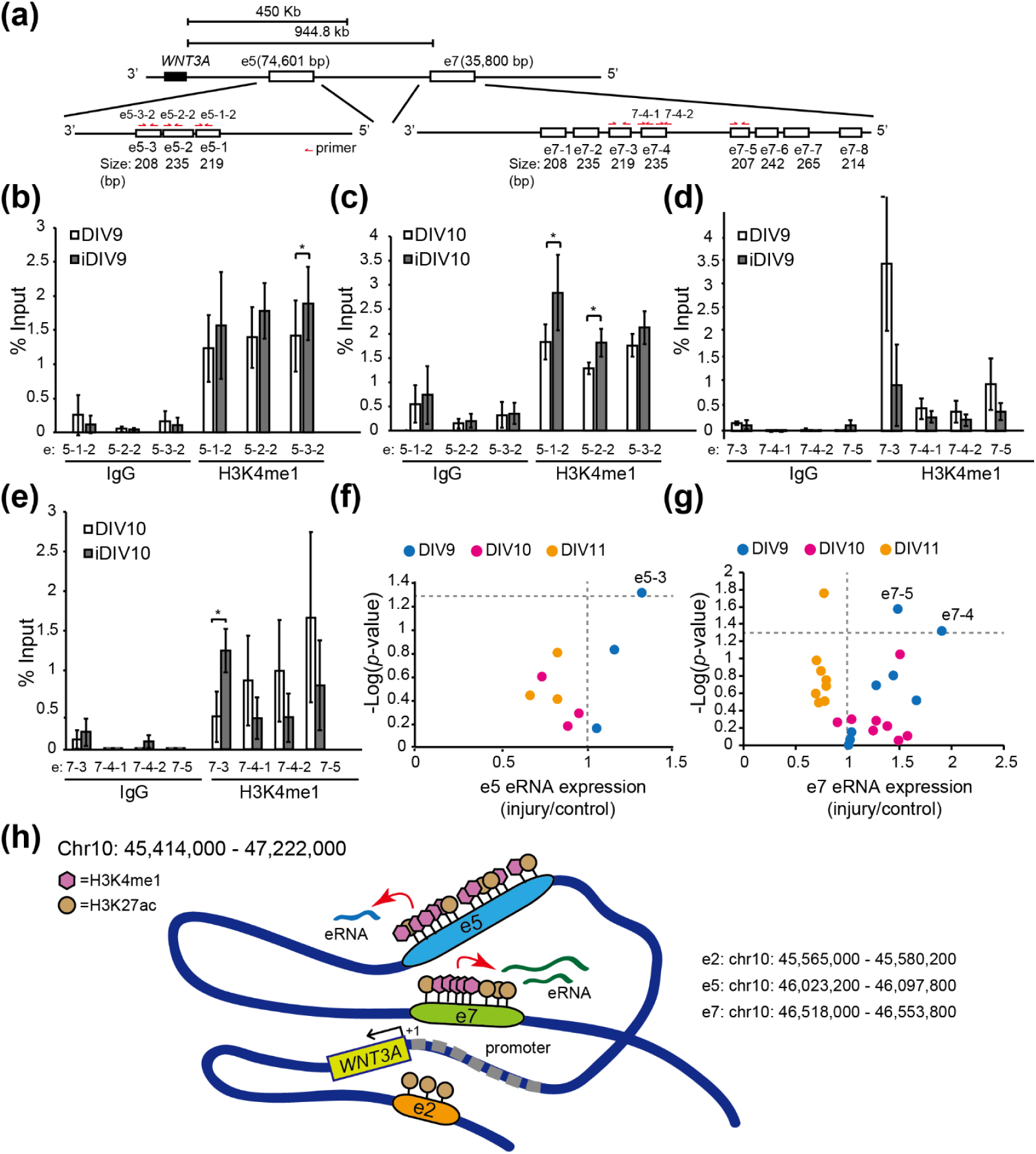
H3K4me1 modifications and eRNAs at putative enhancer regions during neuronal regeneration. (a) Schematic diagram of primer design for ChIP assays at e5 and e7 regions. (b-e) ChIP-qPCR was performed to examine the level of H3K4me1 at the e5 (b-c) and e7 (d-e) regions. IgG enrichment was used as a negative control. Samples were from cortical neurons on DIV9, iDIV9, DIV10 and iDIV10 respectively. Data are mean ± SEM (n = 3 and n = 4 for experiments on DIV9 and DIV10, respectively). \**P* ≤ 0.05 (paired Student’s *t*-test). (f, g) Volcano plots displaying differential expression (DE) of e5 and e7 eRNAs between control and injured samples on DIV9, DIV10 and DIV11, as determined by RT-PCR. The x-axis represents the fold-change of eRNA expression. The y-axis corresponds to –log (*p*-value), determined by paired Student’s *t*-test. Vertical gray dotted line represents the value of DE = 1, while the horizontal gray dotted line borders the cut-off value of –log (*p* = 0.05). eRNA DE values are shown as mean from at least three independent experiments (e5: n = 3; e7: n = 4). (h) Working model of enhancer-regulated *WNT3A* transcription. Dark blue line depicts genomic DNA ranging from predicted enhancers e1 to e10 (chr: 45,414,000-47,222,000). Relative position of the e2, e5, e7 enhancer regions are denoted. Pink hexagon represents H3K4me1. Brown circle represents H3K27ac. eRNAs derived from e5 and e7 are indicated as light blue and green lines, respectively. Red arrows point to the activation of eRNA transcription mediated by increased histone modification during regeneration. Black arrow marks the transcription of *WNT3A*. Promoter region for *WNT3A* is depicted by thick gray dashed line. The TSS is denoted as “+1”.

Together, we propose that the distal enhancer e5 (chr10: 46,023,200-46,097,800) and e7 (chr10: 46,518,000-46,553,800) regions, containing H3K4me1-marked TAD, are spatially brought together to enhance the transcription of eRNAs. These eRNAs may recruit transcriptional components allowing e5/e7 enhancer-promoter looping to coordinate *WNT3A* gene expression. Alternatively, the downstream e2 region (chr10: 45,565,000-45,580,200) loops to interact with promoter of *WNT3A* and induces its expression (Fig. 7h). In the last decade, researchers have gained understanding of the multiple functions of eRNAs in regulating enhancer-promoter looping (Yang *et al*. 2016) and promoter-proximal pausing of RNAPII (Shii *et al*. 2017), which both affect transcription. Therefore, eRNAs produced by active e5/e7 enhancer region may stabilize the long-ranged chromatin looping structure and transcriptional complex to maintain *WNT3A* transcription.

## 4. Discussion

Upon neural injury, various signaling pathways (e.g., STAT3, BMP signaling) are triggered in reactive astrocytes, followed by the production of growth factors (e.g. BDNF, NGF), cytokines (e.g. IL-6, CNTF, CT1), and secreted proteins that are permissive to neuronal regeneration and proliferation of glial cells (Ohtake *et al*. 2016; Anderson *et al*. 2016; Karki *et al*. 2014). The current study characterized the positive effect of WNT3A during regeneration of injured cortical neurons, brain slice and CCI mice model. While there was no previous report showing how *WNT3A* gene may be regulated, this study predicted novel enhancers, e5 (chr10: 46,023,200-46,097,800) and e7 (chr10: 46,518,000-46,553,800) for *WNT3A* gene expression during neuronal regeneration. Histone H3K4me1 modification on this enhancer is likely determine the transcriptional activity of e5/e7 to regulate *WNT3A* expression. Since H3K4me1 was previously shown enriched at active enhancers and to fine-tune transcriptional activity by recruiting chromatin modifiers (Rada-Iglesias 2018), we reasoned that the observed increase of H3K27ac on iDIV10 (ChIP-seq results) could be a consequence of H3K4me1-primed enhancer activation (Fig. 5e). Histone ChIP-qPCR assays demonstrated that H3K4me1 at the e5 region was increased as early as iDIV9, concomitant with increased eRNA transcripts (Fig. 7b, f). Moreover, the variable patterns and limited increase of H3K27ac at the e5 and e7 enhancer on iDIV10 found in the ChIP-seq support a dominant role of H3K4me1 in the transcriptional activity of e5/e7 under the context of neuronal regeneration (Fig. 5e). While H3K4me1 marks primed/active enhancer collaboratively with other histone modifications (such as H3K27ac for active enhancer) and has played a decisive role in enhancer activation, the cellular mechanism leads to transcriptional changes is under-studied (Calo & Wysocka 2013). The enzymatic activity of specific H3K4 methyltransferases, such as the members of the MLL/COMPASS family, MLL3/4, are responsible for H3K4 mono-methylation on enhancers during cell differentiation and the pathogenesis of cancer (Hu *et al*. 2013; Sze & Shilatifard 2016; Lee *et al*. 2013). Augmentation of H3K4me1 on enhancers serves as the binding element for the recruitment of mediators, transcription factors, and other effector proteins that engage in the regulation of chromatin environment and the assembly of transcriptional machinery (Smith & Shilatifard 2010). Recent studies demonstrated that catalytically deficient MLL3/4 and knockout of MLL3/4 in cell lines led to reduction of H3K27ac and target gene expression respectively (Dorighi *et al*. 2017) and that reduced binding of chromatin remodeling complex BAF to enhancer is associated with the depletion of H3K4me1 in mutant mouse embryonic stem cell lines with catalytically inactive MLL3/4 (Local *et al*. 2018). It is possible that e5/e7 enhancer accommodates methyltransferases as well as candidate effector proteins in response to injury for transcriptional activation of *WNT3A* gene. Further characterization of the enzymatic activity and expression level of MLL3/4 and potential transcriptional factor binding at the e5/e7 region during neuronal regeneration would reveal additional regulation.

For therapy, it may be possible to utilize CAQK-conjugated nanoparticles or biomaterial to package modified CRISPR (clustered regularly interspaced short palindromic repeats)/dCas9 constructs that target enhancers of *WNT3A* in injured brain neurons (Dong 2018; Mann *et al*. 2016; Bharadwaj *et al*. 2018). RNA-guided inactive Cas9 protein (dCas9) fused to a SET domain of histone methyl transferase (MLL3/4, a.k.a. KMT2C/2D) may then be delivered to activate the enhancer of *WNT3A*. The feasibility and efficacy of administering nanoparticles containing *WNT3A* enhancer-targeting modified CRISPR constructs may be evaluated and could potentially become a therapeutic strategy for treating brain injury. A major challenge for treating TBI lies in the efficiency of drug delivery and cellular uptake, which is limited by the BBB. In healthy brains, the BBB restricts the influx of small molecules from circulation (Pardridge 2015; Pandey *et al*. 2016). Although WNT3A is not likely to pass through the BBB in an uninjured brain due to its size, the BBB is compromised in response to TBI and may be permissive to WNT3A uptake after intravenous injection (Zhao *et al*. 2016). On the other hand, intranasal delivery of proteins is believed to bypass the BBB in animal models (Zhang *et al*. 2018), and our results demonstrate that this method of WNT3A delivery can produce a promising outcome on regeneration of injured brain neurons. The clinical efficacy of systemic or intranasal administration of protein drugs to the brain of TBI patients remains to be tested (El-Amouri *et al*. 2013; Wei *et al*. 2018). Alternatively, small-molecule activators of WNT3A, such as 6-bromoindirubin-3’-oxime (BIO), may be readily permeable to the BBB and could be considered for clinical use.

## Acknowledgements

We thank Mr. We-Han Chiang for culturing primary cortical neurons for NGS study. We thank Dr. Jin-Wu Tsai from the National Yang-Ming University, Taiwan, for providing technical consultation on brain slice culture. We thank Mr. Hsueh-Chun Chuang and Ms. Hsuan-Hsuan Lee for helping with the brain slice culture. We thank the NGS core facility at the Academia Sinica, Taiwan, for performing the NGS.

## Author contributions

LC and CFK designed the study and performed histone ChIP for ChIP-seq. CYC, JHH, and JL analyzed the ChIP-seq and RNA-seq data. HIC cultured primary neurons, isolated RNAs for RNA-seq and performed qPCR analysis. CYC performed qPCR analysis and histone ChIP experiments. CCW performed brain slice organotypic culture. CCW and PYH conducted animal behavior assays. MZL and SFY performed experiments using CCI animal model and MZL imaged brain slice. KSF performed enhancer RNA and histone ChIP experiments. LC, CYC, MZL, CCW, JHH and CFK wrote the manuscript.

## Data Availability

The data that support the findings of this study are available from the corresponding author upon reasonable request.

